# A conserved motif of *Porphyromonas* Type IX secretion effectors C-terminal secretion signal specifies interactions with the PorKLMN core complex

**DOI:** 10.1101/483123

**Authors:** Maxence S. Vincent, Maïalène Chabalier, Eric Cascales

## Abstract

The Type IX secretion system (T9SS) is a versatile protein transport apparatus restricted to the Bacteroidetes phylum. This multiprotein complex enables secretion of a wide range of effectors, such as protein toxins, filamentous adhesins, enzymes or S-layer subunits. Once translocated in the periplasm through the Sec pathway, recognition and secretion of these cargo proteins rely on a conserved C-terminal domain referred as CTD. However, the precise route followed by the CTD substrates from the periplasm to the cell exterior is yet to be determined. Here we define the interaction network of five different CTDs from the oral pathogen *Porphyromonas gingivalis* with the PorKLMN T9SS trans-envelope complex. We show that these five CTDs interact with the inner membrane-anchored PorM and the PorN periplasmic proteins. We further determine the contribution of the PorM IgG-like periplasmic domains and CTD conserved motifs for PorM-CTD complex formation. These results showed that all five CTDs interact with the core complex in a similar manner, suggesting a conserved mechanism of substrate selection in the periplasm. Our results thus establish a first model of the path followed by the CTD substrates through the T9SS.

## INTRODUCTION

Gram-negative pathogens use secretion systems to deliver effectors and toxins in the external medium or into target cells. Based on the apparatus components and architecture, and on effector recruitment and sorting, secretion systems have been categorized in nine families. The Type IX secretion system (T9SS), that conveys effectors to the milieu or to the cell surface, is restricted to members of *Bacteroidetes* phylum [1,2]. It is now well established that *Bacteroidetes* opportunistic pathogens such as *Porphyromonas gingivalis* and *Tannerela forsythia* employ this multiprotein complex to deliver toxins across the outer membrane [3–5]. Other *Bacteroidetes* members use the T9SS for the secretion of adhesins required for gliding motility, or enzymes required for supplying carbon, heme and metal sources [5–8]. *P. gingivalis,* the main causative agent of periodontal diseases, secretes a plethora of virulence factors at its cell surface, including the gingipains that play an important role in gum infections [9]. The gingipain family comprises three arginine- or lysine-specific cysteine proteinases, RgpB, RgpA and Kgp, which actively promote the degradation of periodontal tissues [9]. More recently, the peptidylarginine deiminase (PPAD), a toxin responsible for protein citrullination and implicated in the development of rheumatoid arthritis, has also been shown to be delivered by the *P. gingivalis* T9SS [10].

While the composition of T9SS varies between species, a number of components are conserved in all the T9SSs identified so far, suggesting the existence of a conserved core [1,2]. This T9SS core comprises three sub-complexes: (i) a trans-envelope complex composed of two inner membrane proteins, PorL and PorM, and of an outer membrane-associated complex composed of the PorK outer membrane lipoprotein and the PorN periplasmic protein, (ii) an outer membrane translocon, and (iii) an attachment complex that comprises the PorU, PorV and PorZ proteins [1,2] (Fig. 1A). Additional conserved components, such a PorP, PorQ, PorT and PorW are yet poorly characterized [1,2]. The PorKLMN proteins assemble onto a > 1.4-MDa trans-envelope complex [5] through a dense network of contacts[11]. The PorL and PorM proteins interact via their transmembrane segments in the inner membrane [11]. The crystal structure of the periplasmic PorM region demonstrated that it forms dimer composed of four domains, called D1-D4, including the D2-D4 Ig-fold-like domains [12]. This domain engages in interaction with the PorKN complex at the outer membrane [11,12]. Electron microscopy (EM) analyses of the purified PorKN complex revealed that it assembles onto 50 nm-diameter ring-like structures [13]. The cryo-EM structure of the SprA translocon (the Sov homologue in *F. johnsioniae*) has been recently solved [14]. It forms a large β-barrel channel crossing the outer membrane and comprising 36 β-strands [14]. The Sov translocon exists in two states: a closed state, in which a plug protein prevents access to the Sov lumen, and an open state, lacking the plug protein but associated with PorV, a protein of the attachment complex [14]. This structure is suggested to be the conformation that will engage the substrate to be translocated [14]. The attachment complex comprises the PorV β-barrel outer membrane protein and the cell surface-exposed PorU and PorZ subunits [15,16]. This attachment complex uses a sortase-like mechanism to proteotically process the C-terminal domain (CTD) of the effectors and to attach the mature effector to anionic LPS [17] (Fig. 1A).

**Figure 1.**
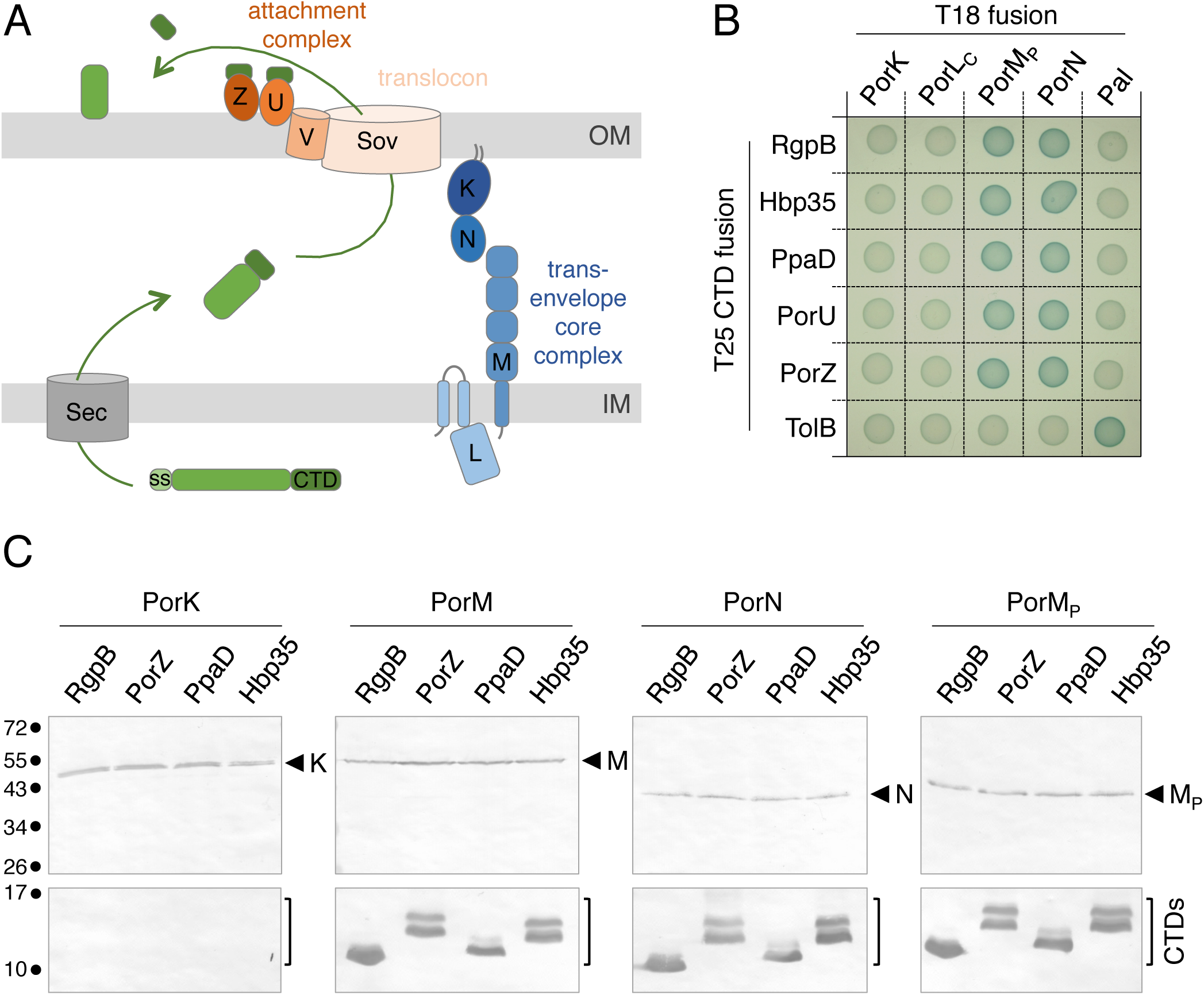
T9SS effector C-terminal domains interact with components of the trans-envelope PorKLMN complex. (A) Schematic representation of the mechanism of T9SS-dependent secretion. Cytoplasmically synthesized effectors (green) are exported through the inner membrane (IM) via the Sec pathway. After cleavage of its N-terminal signal sequence (SS), the C-terminal domain (CTD) targets the effector to the T9SS for translocation across the outer membrane (OM). Once at the cell surface, the CTD is cleaved and the effector is attached to the cell surface or released in the medium. The three main sub-complexes of the T9SS (trans-envelope core complex, blue; translocon, light pink; attachment complex, orange) are shown. The letter indicated the name of the Por protein (e.g., M is PorM). (B) Bacterial two-hybrid analysis. BTH101 reporter cells producing the indicated proteins fused to the T18 and T25 domain of the *Bordetella* adenylate cyclase were spotted on X-Gal-IPTG reporter LB agar plates. The blue color of the colony reports interaction between the two partners. Controls include T18 and T25 fusions to TolB and Pal, two proteins that interact but unrelated to the T9SS. (C) Co-immunoprecipitation assay. Soluble lysates of *E. coli* cells producing periplasmically-targeted VSV-G-tagged CTDs with FLAG-tagged PorK (K), PorM (M), PorN (N) proteins or PorM periplasmic domain (PorM_P_, M_P_) were subjected to immunoprecipitation with anti-FLAG-coupled beads. The immunoprecipitated material were separated by 15% acrylamide SDS-PAGE and immunodetected with anti-FLAG (upper panel) and anti-VSV-G (lower panel) monoclonal antibodies. The position of the CTDs and Por proteins or domains are indicated on the right of each panel. Molecular weight markers (in kDa) are indicated on left

Bioinformatics analyses showed that the T9SS effectors CTD is a 60-110 C-terminal extension that is found in > 650 proteins produced by Bacteroidetes T9SS^+^ species but is absent in proteins from *Bacteroidetes* T9SS- species, or in non-*Bacteroidetes* species [18]. Deletion of the CTD leads to the accumulation of the cargo protein in the periplasm [19,20], which is in agreement with the presence of a signal peptide at the N-terminus of T9SS effectors (Fig. 1A). The CTD has been shown to be necessary and sufficient to promote T9SS-dependent translocation of reporter proteins such as the GFP [21–23]. It is thus commonly accepted that the CTD corresponds to the secretion signal that is recognized by the T9SS to select proteins to be translocated. The CTD extension is therefore present on all the T9SS substrates, including the cell surface-exposed PorU and PorZ proteins of the attachment complex [17,24,25]. However, in the cases of PorU and PorZ, the CTD is not cleaved after transport, suggesting differences during the final maturation of CTD-containing substrates [17,24,25]. The atomic structures of four *P. gingivalis* CTDs have been recently solved, showing a conserved seven antiparallel β-strands β-sandwich Ig-like fold [10,25–27]. Interestingly, sequence alignment of *Bacteroidetes* CTDs defined five motifs (A-E) including three conserved motifs (B, D and E; [18,19]). Deletion of the last 22 amino-acids of the Hbp35 CTD, a segment that comprises the D and E motifs, showed that these two motifs are necessary for efficient secretion of the effectors [21], and may participate to the proper folding of the CTD or to the recognition of the T9SS.

While genomic and structural information have been accumulated on T9SS effector CTDs in the recent years, we still lack a functional understanding on how CTD effectors engage with the T9SS. Here, we provide evidence that the CTDs from five T9SS substrates, RgpB, Hbp35, PpaD, PorU and PorZ, interact with two subunits of the trans-envelope complex, PorM and PorN. We then show that CTDs engage in at least two different interactions with the D2-D3 and D4 periplasmic regions of PorM. Finally, mutagenesis of the CTDs B, D and E motifs coupled to interaction studies demonstrate that motif D is required for complex formation with PorM. We then propose a model on the initial steps of T9SS effectors recruitment to the T9SS, prior to their engagement into the T9SS translocon.

## RESULTS

### T9SS effector C-terminal domain interacts with PorM and PorN

In the current model, T9SS substrates are selected in the periplasm and addressed to the SprA/Sov translocon for their translocation across the outer membrane (Fig. 1A; [1,2]). How the substrate CTDs are recognized by the T9SS is yet unknown. To address this question, we tested whether the CTDs from various *P. gingivalis* T9SS substrates interact with the PorKLMN subunits, which form the conserved trans-envelope complex of the apparatus [11]. The CTDs used in this study include that of the RgpB arginine-specific cysteine proteinase (PGN_1466; amino-acids 661-736), the Hbp35 hemin-binding protein (PGN_0659; amino- acids 266-344), the PpaD peptidyl-arginine deiminase (PGN_0898; amino-acids 478-557), and the PorU (PGN_0022; amino-acids 1059-1158) and PorZ (PGN_0509; amino-acids 680- 776) attachment complex proteins. The sequences encoding the CTDs were cloned into the pKT25 bacterial two-hybrid (BACTH) vector, downstream the T25 domain sequence of the *Bordetella* adenylate cyclase. These T25-CTD fusion proteins were tested against T18 constructs fused to the soluble domains of the PorK (lacking the signal peptide and N-terminal acylated cysteine residue), PorL (cytoplasmic domain), PorM (periplasmic domain), and PorN (lacking the signal peptide) proteins [11]. The results shown on X-Gal reporter plates demonstrate that all five CTDs interact with the PorM periplasmic domain and the PorN periplasmic protein (Fig. 1B). No interaction was detected with PorK or with the PorL cytoplasmic domain. These interactions were confirmed by co-immunoprecipitation (co-IP) in the heterologous host *E. coli* K-12. VSV-G-tagged CTDs were fused to a signal sequence for periplasmic targeting, and tested for their interaction with full-length FLAG-tagged PorK, PorM and PorN proteins. Because PorL has a very limited 8-amino-acid periplasmic loop (see Fig. 1A), it was not included in the study. Fig. 1C shows that the CTDs specifically co-immunoprecipitate with PorM and PorN. In addition, the PorM periplasmic domain, exported to the periplasm, also interacts with the CTDs (Fig. 1C). As shown for the BACTH assay, the CTDs do not co-precipitate with the PorK lipoprotein. Because of the absence of the other T9SS genes in *E. coli* K-12, we conclude that the CTDs directly interact with both the PorM periplasmic domain and PorN.

### T9SS effector CTDs interact with the Ig-fold periplasmic domains of PorM

The recently solved crystal structures of the periplasmic fragments of the *P. gingivalis* PorM (PDB: 6EY0 and 6EY5) and its GldM homologue in *F. johnsionae* (PDB: 6EY4) showed that they comprise 4 domains: an N-terminal all α-helical domain (D1) followed by three Ig-likeβ-domains (D2, D3 and D4) (Fig. 2A; [12]). We used BACTH and co-immunoprecipitation assays to test the contribution of these different domains for CTDs interaction. The results shown in Fig. 2B and 2C demonstrate that the CTDs interact with the D2-D3-D4 Ig-like domains. To further delineate the site of contact of the CTD, we engineered constructs producing the D2-D3 and D4 domains. Interestingly, protein-protein interaction analyses show that the CTDs make contacts with both the D2-D3 and D4 constructs (Fig. 2B and 2C) suggesting that the CTDs engage in interaction with two regions of PorM concomitantly or subsequently.

**Figure 2.**
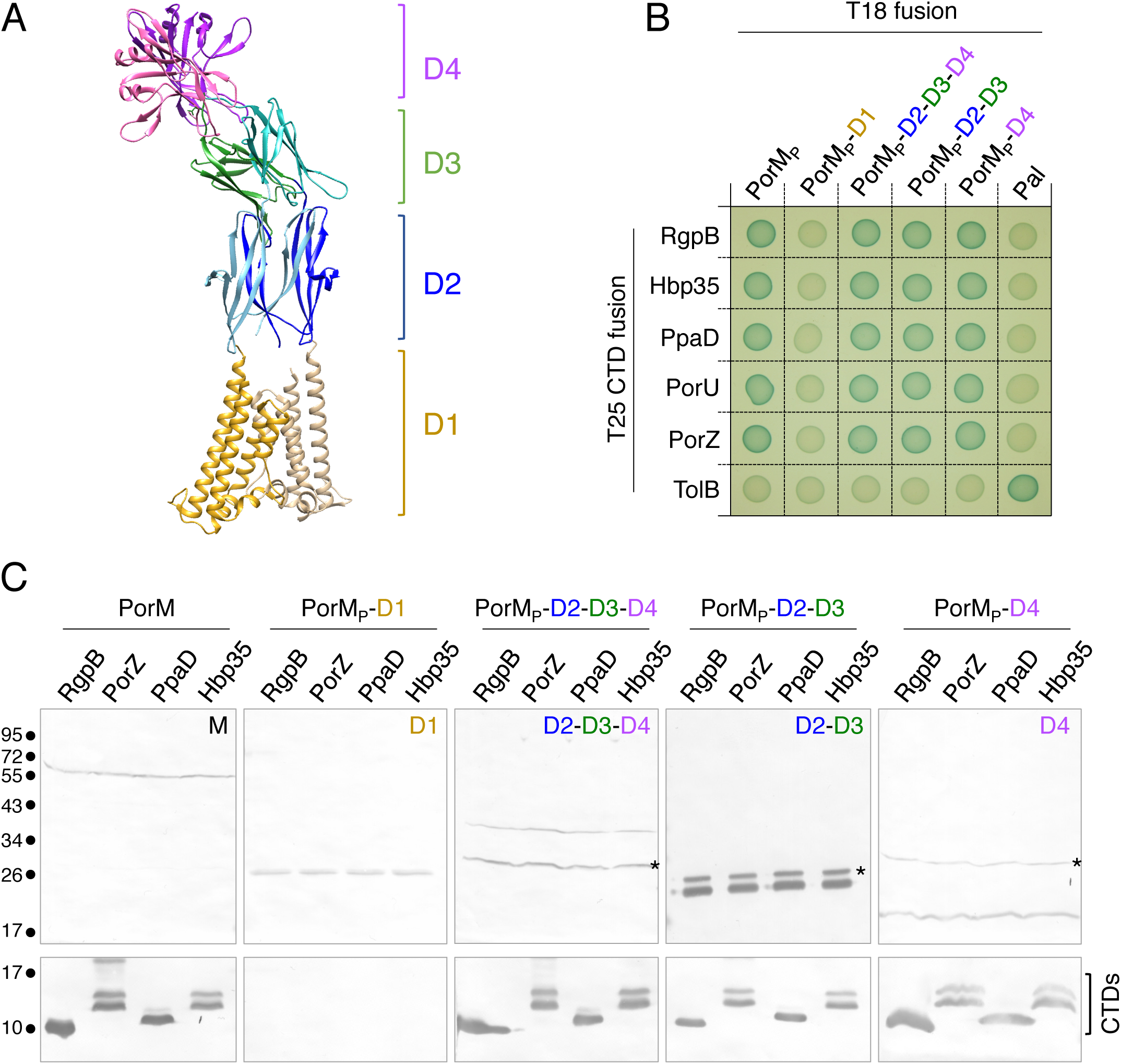
T9SS effector CTDs interact with the PorM Ig-like periplasmic domains. (A) Ribbon representation of the structure of the dimeric PorM periplasmic region (PDB: D1: 6EY0, D2-D4: 6EY5; [12]). The different domains are represented with a color code (gold, D1; blue, D2; green, D3; pink, D4). (B) Bacterial two-hybrid analysis. BTH101 reporter cells producing the indicated domains fused to the T18 and T25 domain of the *Bordetella* adenylate cyclase were spotted on X-Gal-IPTG reporter LB agar plates. The blue color of the colony reports interaction between the two partners. Controls include T18 and T25 fusions to TolB and Pal, two proteins that interact but unrelated to the T9SS. (C) Soluble lysates of *E. coli* cells producing periplasmically-targeted VSV-G-tagged CTDs with FLAG-tagged PorM (M), or PorM domains (D1, D2-D3- D4, D2-D3 or D4) were subjected to immunoprecipitation with anti-FLAG-coupled beads. The immunoprecipitated material were separated by 15% acrylamide SDS-PAGE and immunodetected with anti- FLAG (upper panel) and anti-VSV-G (lower panel) monoclonal antibodies. The asteriks indicate the position of anti-FLAG antibody light chain. Molecular weight markers (in kDa) are indicated on left.

### A CTD conserved motif mediates interactions with PorM

Sequence analyses of the T9SS effector CTDs previously showed that they comprise three conserved motifs, referred as motifs B, D and E [18,19]. A sequence alignment of the five CTDs used in this study readily identified these three motifs (Supplemental Fig. 1A, Fig. 3A). Furthermore, a comparison of the available structures of these CTDs (RgpB [PDB: 5HFS; [26]], HbpB35 [PDB: 5Y1A;[27]], PorZ [PDB: 5M11;[25]], and PpaD [PDB: 4YT9; [10]])shows that these structures are superimposable, and hence that the B, D and E motifs share conserved locations (Supplemental Fig. 1B). To better understand the role of these motifs in CTD selection, we engineered substitution variants in which the invariable or most conserved residues of each motif were mutated (Fig. 3A). These variants were then tested for their ability to interact with PorM and PorN. Whereas none of the motifs is involved in the interaction with PorN, mutation of motif D resulted in the disruption of the interaction with PorM (Fig. 3B and 3C).

**Figure 3.**
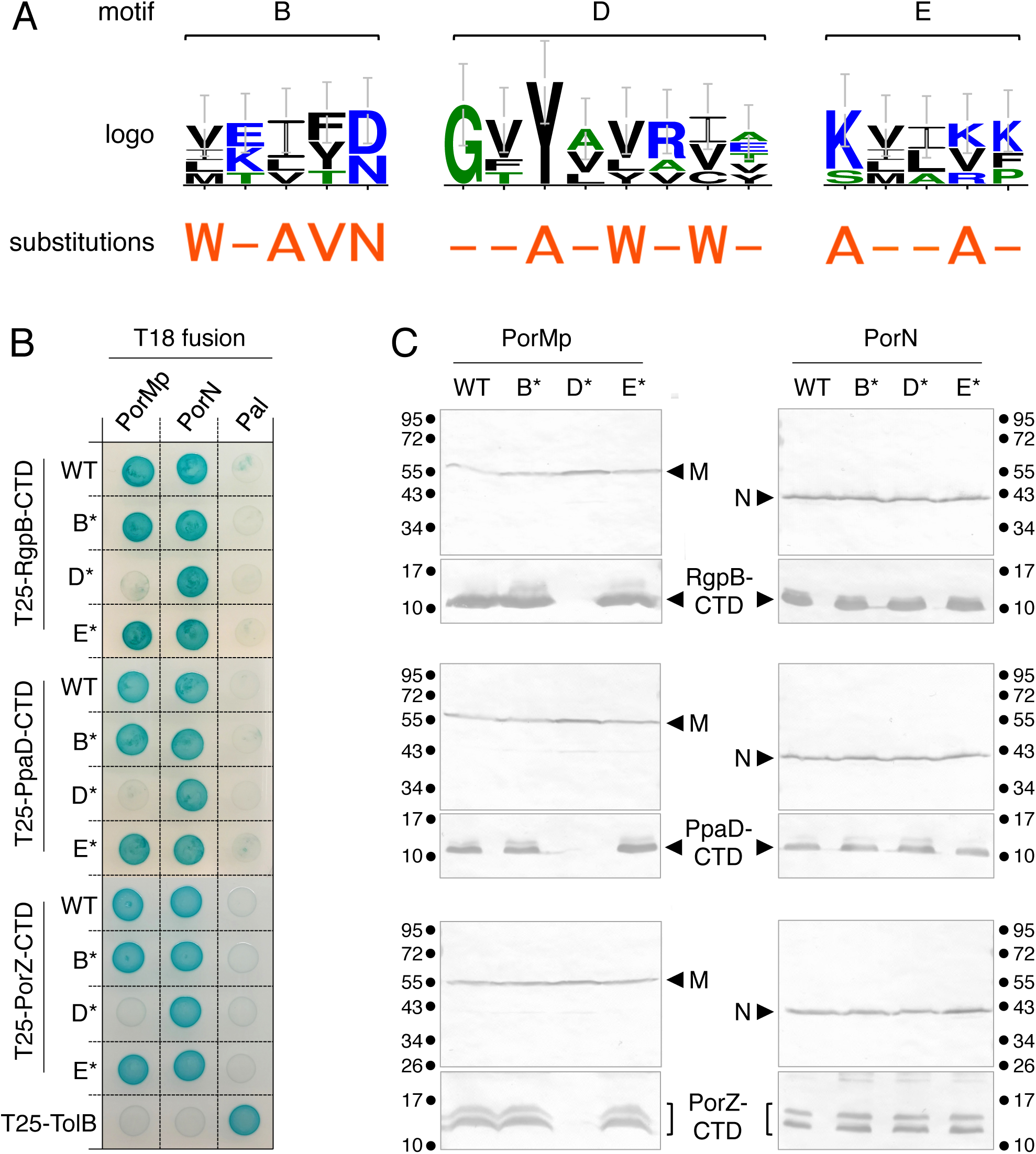
Contribution of CTD conserved motifs to interaction with PorM and PorN. (A) Logo representations of the conserved B, D and E motifs based on a sequence alignment of the 5 CTDs used in this study. For each position, the height of each amino-acid one-letter code represents its frequency at this position in the CTD effectors. The substitutions engineered in each of these motifs are indicated below, in orange. (B) Bacterial two-hybrid analysis. BTH101 reporter cells producing the indicated CTD variants (B*, D* and E* corresponds to the motifs B, D and E variants) fused to the T18 and T25 domain of the *Bordetella* adenylate cyclase were spotted on X-Gal-IPTG reporter LB agar plates. The blue color of the colony reports interaction between the two partners. Controls include T18 and T25 fusions to TolB and Pal, two proteins that interact but unrelated to the T9SS. (C) Soluble lysates of *E. coli* cells producing periplasmically-targeted VSV-G-tagged CTD variants with FLAG-tagged PorM periplasmic domain (M_P_, left panels), or PorN (right panels) were subjected to immunoprecipitation with anti-FLAG-coupled beads. The immunoprecipitated material were separated by 15% acrylamide SDS-PAGE and immunodetected with anti-FLAG (upper panel) and anti-VSV-G (lower panel) monoclonal antibodies. Molecular weight markers (in kDa) are indicated.

## DISCUSSION

In this study, we provide the first evidence of effector CTD interactions with conserved core components of the T9SS. We show that the CTDs from five different T9SS effectors interact with the periplasmic domain of the PorM inner membrane protein and the periplasmic PorN protein. We further show that the CTDs interact with two regions with the PorM periplasmic region: the D2-D3 and D4 domains. We then demonstrate that a conserved motif within T9SS CTDs, motif D, is required for proper interaction with PorM.

During the secretion process, T9SS effectors are first exported to the periplasm via the Sec pathway. They then have to engage with the T9SS apparatus, in the periplasm, to be translocated across the outer membrane. Our results, showing interactions with the periplasmic region of PorM and the PorN periplasmic protein, therefore suggest that these contacts represent the early steps of T9SS-dependent effector translocation. PorM and PorN may therefore constitute the initial checkpoint to verify and select CTD proteins to be translocated. Indeed, the observation that the CTD motif D is required for PorM binding suggests that it may correspond to a selection motif necessary for engagement of the substrate in the type IX pathway. Interestingly, all CTDs tested, which represent diverse families (effectors with different activities, subunits of the attachment complex), interact with the same components of the trans-envelope complex. In addition, the motif D of the three T9SS substrates tested, RgpB, PpaD and PorZ, is required for complex formation with PorM (Fig. 4A and B). These results suggest that CTDs specify engagement of the cargo proteins into a conserved transport pathway, or at least that the initial step of T9SS substrates selection in the periplasm is conserved. Indeed, studies performed in *Flavobacterium johnsioniae* showed that translocation of various CTD substrates differ in their requirement for the PorU and PorV attachment complex proteins [28], suggesting that later stages of CTD transport may require specific signals that are unlikely to be conserved across all T9SS substrate CTDs.

**Figure 4.**
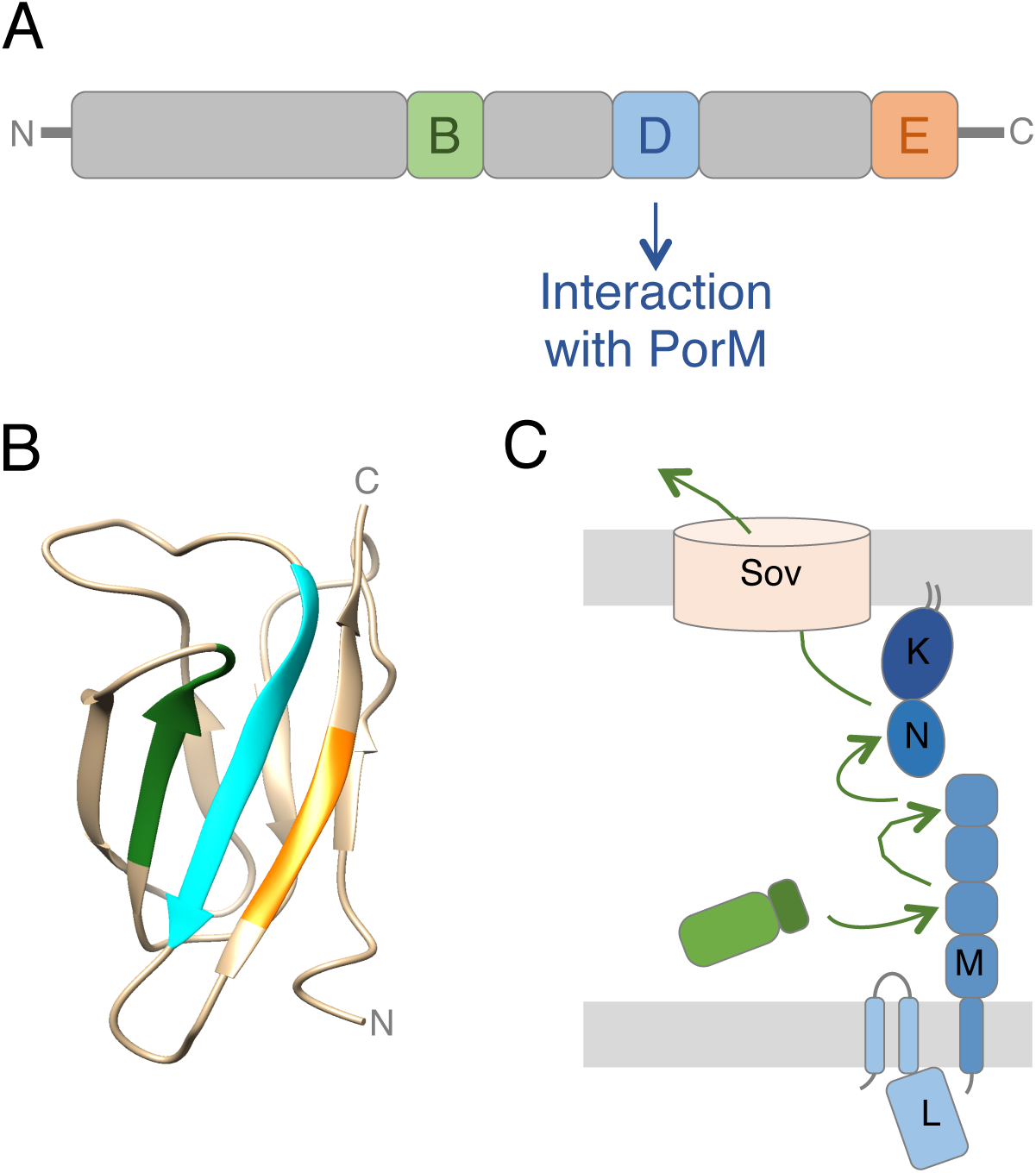
CTD architecture, structure and transport. (A) Schematic representation ofT9SS effector architecture. The positions of the conserved B (green), D (blue), and E (orange) motifs are indicated, as well as their contribution to the interaction with T9SS components. (B) Structure of the RgpB CTD. The positions of the B (green), D (blue), and E (orange) motifs are highlighted. (C) Proposed mechanism of transport of effectors by the T9SS. Once in the periplasm, the CTD sequentially interacts with the PorM D2, D3 and D4 domains, and PorN to be addressed to the Sov translocon.

Our study did not reveal a specific conserved motif necessary for the interaction with PorN. One may hypothesize that this interaction requires one of the less conserved motifs not tested in this study (A or C), a combination of motifs, or additional residues that are not part of a linear motifs but close in space. Comparison of the available CTD tri-dimensional structures may reveal conserved structural features. On the other hand, we did not specify the role of motifs B and E. We suggest that these motifs could be important for the proper folding of the CTD or for mediating contacts with other T9SS subunits such as the translocon or the attachment complex. Motif B comprises hydrophobic side chains and aromatic rings that protrude towards the interior of the β-sandwich and therefore likely involved in the stability of the CTD structure. It would be interesting to test CTD interaction with other T9SS subunits and define the contribution of the conserved motifs in these interactions.

A striking observation of our study is that the CTDs interact with different regions of the PorM periplasmic fragment, domains D2-D3 and D4. Interestingly, mutations within motif D abolish CTDs binding to PorM, suggesting that a single motif is engaged in the interaction with the D2-D3 and D4 domains. These interactions could be either concomitant or sequential. Because the structure of the PorM periplasmic fragment presents an elongated structure (see Fig. 2A; [12], we favour the second hypothesis. However, the crystal structures of PorM and of its *F. johnsioniae* GldM homologue showed that a large bend occurs in the PorM structure compared to that of GldM [12]. Based on this difference, we hypothesized that these two structures may represent two different states of the PorM/GldM conformation during the secretion process. If this turns to be true, one can imagine that this bending brings the D2-D3 and D4 domains at close proximity to accommodate the CTD in a horseshoe-like structure. On the other hand, it is interesting to note that the PorM/GldM D2, D3 and D4 domains share a conserved Ig-like fold. Therefore, the CTD motif D may engage in sequential interactions with these different domains by using the same type of interactions. Interestingly, while the Ig-like fold is conserved, there are significant differences between these three domains, notably regarding the organisation of the secondary structures. One exciting hypothesis is that these differences contribute to different affinities to the CTDs. By sequentially interacting with the PorM D2, D3 and D4 domains, the CTDs will be shuttled to the outer membrane-associated PorKN complex or to the Sov translocon.

Based on our data, we propose a working model regarding effector selection and transport by the T9SS (Fig. 4C). Newly synthesized effectors are exported to the periplasm via the Sec pathway in an unfolded state. Although effector folding in the periplasm has not been evidenced, the requirement for a specific Ig-like domain for T9SS-dependent transport, the internal diameter of the outer membrane translocon [14], and the observation that T9SS substrates sequestered in the periplasm are enzymatically active [19,29], we anticipate that effector folding occurs in the periplasm before transport by the Type IX secretion apparatus. In our model, the PorM D2 domain selects periplasmic T9SS effectors by recognition of CTD motif D. Increasing affinities of the CTD for the D3 and D4 domains pilot the effector close to the outer membrane where it engages into the translocon for its translocation to the cell surface. While many of these steps need to be experimentally tested, this model paves the way for a better understanding of effector translocation by the T9SS.

## MATERIAL AND METHODS

### Bacterial strains, medium, and growth conditions

Strains used in this study are listed in Table S1. *Porphyromonas gingivalis* ATCC33277 was used as source of DNA for cloning. *P. gingivalis* was grown anaerobically in brain-heart infusion medium supplemented with menadiol (0.5 μg.mL^-1^) and hemin (5 μg.mL^-1^). *Escherichia coli* DH5α (New England Biolabs), W3110 (laboratory collection) and BTH101 [30] were used for cloning procedures, co-immunoprecipitation, and bacterial two-hybrid assays, respectively. *E. coli* cells were routinely grown aerobically in Lysogeny broth (LB) at 30°C or 37°C. Plasmids were maintained by addition of ampicillin (100 μg.mL^-1^), or kanamycin (50 μg.m^-1^). When necessary, gene expression was induced with 0.1-0.5 mM isopropyl-β-thio-galactoside (IPTG) or 0.05 μg.mL^-1^ of anhydrotetracycline (AHT).

### Plasmid construction

Plasmids used for this study are listed in Table S1. BACTH and pASK-IBA vectors encoding the PorKLMN proteins or domains have been previously described [11,12,31]. DNA sequences corresponding to the T9SS effector CTDs were amplified from *P. gingivalis* genomic DNA extracted from 6×10^9^ cells using a DNA purification kit (DNeasy Blood & Tissue, Qiagen). PCRs were performed with a Biometra thermocycler using Q^®^ DNA polymerase (New England Biolabs) and custom oligonucleotides synthesized by Sigma-Aldrich (listed in Table S1). BACTH vectors encoding T25-RgpB and T25-Hbp35 CTDs were constructed by standard restriction/ligation cloning. PCR products bearing 5′ XbaI and 3′ KpnI sites were digested by the corresponding restriction enzymes (New England Biolabs) and inserted into the pKT25 (fusion at the C terminus of the T25 domain)vector [30] digested with the same enzymes. All other plasmids were engineered by restriction-free cloning [32] as described previously [33]. Briefly, the DNA fragment of interest was amplified using oligonucleotides introducing extensions annealing to the target vector. The double-stranded product of the first PCR has then been used as oligonucleotides for a second PCR using the target vector (pASK- IBA4, pKT25) as template. PCR products were then treated with DpnI to eliminate template plasmids and transformed into DH5α-competent cells. Substitutions of the CTD motifs were introduced by quick-change site-directed mutagenesis using complementary pairs of oligonucleotides bearing the desired base substitutions and the Pfu Turbo high fidelity polymerase (Agilent technologies). PCR products were then treated with DpnI to eliminate template plasmids and transformed into DH5α- competent cells. All constructs were verified by colony PCR and DNA sequencing (Eurofins sequencing).

### Protein-protein interaction assays

#### Bacterial two-hybrid

*Bordetella* adenylate cyclase-based bacterial two-hybrid assays [30] were performed as previously described [34]. Briefly, the proteins to be tested were fused to the isolated T18 and T25 catalytic domains of the *Bordetella* adenylate cyclase. After introduction of the two plasmids producing the fusion proteins into the reporter BTH101 strain, plates were incubated at 30 °C for 48 h. Three independent colonies for each transformation were inoculated into 600 μl of LB medium supplemented with ampicillin, kanamycin, and IPTG (0.5 mM). After overnight growth at 30°C, 10 μl of each culture was dropped onto LB plates supplemented with ampicillin, kanamycin, IPTG, and 40 μg.mL^-1^ of 5-bromo-4-chloro-3-indolyl β-D-galactopyranoside (X-gal, Euromedex) and incubated for 6-16 h at 30°C. Controls included interaction assays with TolB/Pal, a protein pairs unrelated to the T9SS. The experiments were done at least in triplicate, and a representative result is shown.

#### Co-immunoprecipitation

Co-immunoprecipitation assays were performed as previously described [35] with modifications. W3110 cells producing the protein of interest were grown to an Optical density at 600 nm (*A*_600_) of ~0.4, and the expression of the cloned genes was induced with 0.5 μg.mL^-1^ of AHT for 0.75-1 h. Then, 10^10^cells were harvested, and the pellets were resuspended in 1 mL of LyticB buffer (Sigma-Aldrich) supplemented with 100 μg.mL^-1^lysozyme, 100 μg.mL^-1^ DNase, protease inhibitors (Complete, Roche), and 0.4 mM octylphenoxy poly(ethyleneoxy)ethanol (Igepal^®^ CA-630, Sigma-Aldrich). After incubation for 20 min at 25°C, lysates were clarified by centrifugation at 20,000×g for 10 min. Cell lysates corresponding to the interaction to be tested were mixed and incubated for 30 min on a wheel, and the mixture was applied on anti-FLAG M2 affinity gel (Sigma-Aldrich). After 2 h of incubation, the beads were washed three times with 1 mL of 20 mM Tris–HCl (pH 7.5) and 100 mM NaCl, resuspended in 25 μL of Laemmli loading buffer, boiled for 10 min, and subjected to sodium-dodecyl-sulfate poly-acrylamide gel electrophoresis (SDS-PAGE) and immunodetection analyses.

### Miscellaneous

SDS-PAGE and protein transfer were performed with mini-Protean apparatuses (Biorad) using standard protocols. For immunostaining, proteins were transferred onto nitrocellulose membranes, and immunoblots were probed with primary antibodies and goat secondary antibodies coupled to alkaline phosphatase and were developed in alkaline buffer in presence of 5-bromo-4- chloro-3-indolylphosphate and nitroblue tetrazolium. The anti-FLAG (clone M2, Sigma-Aldrich), and anti-VSV-G (clone P5D4, Sigma-Aldrich) monoclonal antibodies, and alkaline phosphatase-conjugated goat anti-mouse secondary antibodies (Beckman Coulter) have been purchased as indicated and used as recommended by the manufacturer

## Supporting information

## ACKNOWLEDGEMENTS

We thank the members of the Cascales, Lloubès, Cambillau/Roussel, Bouveret and Sturgis research groups for insightful discussions, Wanassa Beroual for help with the constructions of the PpaD CTD- encoding vectors, and Moly Ba, Isabelle Bringer, Annick Brun and Olivier Uderso for technical assistance. This work was supported by the Aix-Marseille Université (AMU), the Centre National de la Recherche Scientifique (CNRS), and grants from the Agence Nationale de la Recherche (ANR-15-CE11-0039-01) and from the Excellence Initiative of Aix-Marseille University (A*MIDEX, A-M- AAP-ID-17-33-170301-07.22), a French “Investissements d’Avenir” programme. M.S.V. was supported by a doctoral fellowship from the French Ministère de la Recherche, and an end-of-thesis fellowship from the Fondation pour la Recherche Médicale (FDT2018-05005242).

