## Supplementary material for "A conserved motif of *Porphyromonas* Type IX secretion effectors C-terminal secretion signal specifies interactions with the PorKLMN core complex"

**SUPPLEMENTAL TABLE S1. Strains, plasmids and oligonucleotides used in this study.**

### STRAINS

| Strains | Description and genotype | Source |
|---|---|---|
| *Porphyromonas gingivalis* | | |
| DSM20709 | WT *Porphyromonas gingivalis* (ATCC33277/DSM20709) | DSMZ collection |
| *Escherichia coli* K12 | | |
| DH5α | F-, Δ(*argF*-lac)U169, *phoA*, *supE44*, Δ(*lacZ*)M15, *relA*, *endA*, *thi*, *hsdR* | Laboratory collection |
| W3110 | F-, lambda-IN(*rrnD-rrnE*)1 *rph*-1 | Laboratory collection |
| BTH101 | F-, *cya99, araD139, galE15, gakK16, rpsL, hsdR, mcrAB* | Karimova et al., 1998 |

### PLASMIDS

| Plasmid | Description and main characteristics | Source |
|---|---|---|
| Expression vectors | | |
| pASK-IBA37(+) | Expression vector, AHT-inducible, $Amp^R$ | IBA technology |
| pIBA-$PorM_{FLAG}$ | *P. gingivalis porM* gene cloned into pASK-IBA37(+), N-terminal FLAG epitope | Vincent et al., 2017 |
| pIBA-$PorN_{FLAG}$ | *P. gingivalis porN* gene cloned into pASK-IBA37(+), C-terminal FLAG epitope | Vincent et al., 2017 |
| pASK-IBA4 | Expression vector, AHT-inducible, OmpA signal sequence, $Amp^R$ | IBA technology |
| pIBA4-$PorK_{FLAG}$ | *P. gingivalis porK* without signal sequence cloned into pASK-IBA4, C-terminal FLAG epitope | Vincent et al., 2017 |
| pIBA-$PorM_{PFLAG}$ | *P. gingivalis porM* periplasmic domain (aa 36-516) cloned into pASK-IBA4, N-terminal FLAG epitope | Vincent et al., 2017 |

| | | |
|---|---|---|
| pIBA-PorM$_P$-D1$_{FLAG}$ | *P. gingivalis porM* D1 domain (aa 44-217) cloned into pASK-IBA4, N-terminal FLAG epitope | Leone et al., 2018 |
| pIBA-PorM$_P$-D2-D3-D4$_{FLAG}$ | *P. gingivalis porM* D2-D3-D4 domains (aa 225-516) cloned into pASK-IBA4, N-terminal FLAG epitope | Leone et al., 2018 |
| pIBA-PorM$_P$-D2-D3$_{FLAG}$ | *P. gingivalis porM* D2-D3 domains (aa 225-399) cloned into pASK-IBA4, N-terminal FLAG epitope | Leone et al., 2018 |
| pIBA-PorM$_P$-D4$_{FLAG}$ | *P. gingivalis porM* D4 domains (aa 400-516) cloned into pASK-IBA4, N-terminal FLAG epitope | Leone et al., 2018 |
| pIBA-RgpB$_V$ | *P. gingivalis rgpB* CTD (aa 661-736) cloned into pASK-IBA4, N-terminal VSV-G epitope | This study |
| pIBA-Hbp35$_V$ | *P. gingivalis hbp35* CTD (aa 226-344) cloned into pASK-IBA4, N-terminal VSV-G epitope | This study |
| pIBA-PpaD$_V$ | *P. gingivalis ppaD* CTD (aa 478-557) cloned into pASK-IBA4, N-terminal VSV-G epitope | This study |
| pIBA-PorU$_V$ | *P. gingivalis porU* CTD (aa 1059-1158) cloned into pASK-IBA4, N-terminal VSV-G epitope | This study |
| pIBA-PorZ$_V$ | *P. gingivalis porZ* CTD (aa 680-776) cloned into pASK-IBA4, N-terminal VSV-G epitope | This study |
| pIBA-RgpB$_V$-B* | Leu692Trp-Ile694Ala-Phe695Val-Asp696Asn substitutions in pIBA-RgpB$_V$ | This study |
| pIBA-RgpB$_V$-D* | Tyr720Ala-Val722Trp-Ile724Trp substitutions in pIBA-RgpB$_V$ | This study |
| pIBA-RgpB$_V$-E* | Lys732Ala-Lys736Ala substitutions in pIBA-RgpB$_V$ | This study |
| pIBA-PpaD$_V$-B* | Ile504Trp-Leu506Ala-Tyr507Val-Asn508Asp substitutions in pIBA-PpaD$_V$ | This study |
| pIBA-PpaD$_V$-D* | Tyr540Ala-Leu522Trp-Val544Trp substitutions in pIBA-PpaD$_V$ | This study |
| pIBA-PpaD$_V$-E* | Lys554Ala-Lys557Ala substitutions in pIBA-PpaD$_V$ | This study |
| pIBA-PorZ$_V$-B* | Val718Trp-Ile720Ala-Thr721Val-Asp722Asn substitutions in pIBA-PorZ$_V$ | This study |
| pIBA-PorZ$_V$-D* | Tyr755Ala-Val757Trp-Val759Trp substitutions in pIBA-PorZ$_V$ | This study |
| pIBA-PorZ$_V$-E* | Lys768Ala-Arg771Ala substitutions in pIBA-PorZ$_V$ | This study |

Bacterial Two-Hybrid (BACTH)

| | | |
|---|---|---|
| pUT18C | BACTH vector, ColE1 origin, *P*lac, T18 domain of *Bordetella* adenylate cyclase, Amp$^R$, | Karimova et al., 1998 |
| pKT25 | BACTH vector, P15A origin, *P*lac, T25 domain of *Bordetella* adenylate cyclase, Kan$^R$ | Karimova et al., 1998 |
| pT18-PorK | *P. gingivalis porK* without signal sequence, cloned downstream T18 into pUT18C | Vincent et al., 2017 |
| pT18-PorL$_C$ | *P. gingivalis porL* cytoplasmic domain (amino-acids 73-309), cloned downstream T18 into pUT18C | Vincent et al., 2016 |
| pT18-PorM | *P. gingivalis porM* cloned downstream T18 into pUT18C | Vincent et al., 2017 |
| pT18-PorM$_P$ | *P. gingivalis porM* periplasmic domain (amino-acids 36-516) cloned downstream T18 into pUT18C | Vincent et al., 2017 |
| pT18-PorN | *P. gingivalis porN* without signal sequence, cloned downstream T18 into pUT18C | Vincent et al., 2017 |
| pT18-PorM$_P$-D1 | *P. gingivalis porM* D1 domain (aa 44-217) cloned downstream T18 into pUT18C | Leone et al., 2018 |
| pT18-PorM$_P$-D2-D3-D4 | *P. gingivalis porM* D2-D3-D4 domains (aa 225-516) cloned downstream T18 into pUT18C | Leone et al., 2018 |
| pT18-PorM$_P$-D2-D3 | *P. gingivalis porM* D2-D3 domains (aa 225-399) cloned downstream T18 into pUT18C | Leone et al., 2018 |
| pT18-PorM$_P$-D4 | *P. gingivalis porM* D4 domains (aa 400-516) cloned downstream T18 into pUT18C | Leone et al., 2018 |
| pT18-Pal | *E. coli pal* without signal sequence, cloned downstream T18 into pUT18C | Battesti and Bouveret, 2007 |

| | | |
|---|---|---|
| pT25-RgpB | *P. gingivalis rgpB* CTD (aa 661-736) cloned downstream T25 into pKTN25 | This study |
| pT25-Hbp35 | *P. gingivalis hbp35* CTD (aa 226-344) cloned downstream T25 into pKTN25 | This study |
| pT25-PpaD | *P. gingivalis ppaD* CTD (aa 478-557) cloned downstream T25 into pKTN25 | This study |
| pT25-PorU | *P. gingivalis porU* CTD (aa 1059-1158) cloned downstream T25 into pKTN25 | This study |
| pT25-PorZ | *P. gingivalis porZ* CTD (aa 680-776) cloned downstream T25 into pKTN25 | This study |
| pT25-RgpB$_V$-B* | Leu692Trp-Ile694Ala-Phe695Val-Asp696Asn substitutions in pT25-RgpB$_V$ | This study |
| pT25-RgpB$_V$-D* | Tyr720Ala-Val722Trp-Ile724Trp substitutions in pT25-RgpB$_V$ | This study |
| pT25-RgpB$_V$-E* | Lys732Ala-Lys736Ala substitutions in pT25-RgpB$_V$ | This study |
| pT25-PpaD$_V$-B* | Ile504Trp-Leu506Ala-Tyr507Val-Asn508Asp substitutions in pT25-PpaD$_V$ | This study |
| pT25-PpaD$_V$-D* | Tyr540Ala-Leu522Trp-Val544Trp substitutions in pT25-PpaD$_V$ | This study |
| pT25-PpaD$_V$-E* | Lys554Ala-Lys557Ala substitutions in pT25-PpaD$_V$ | This study |
| pT25-PorZ$_V$-B* | Val718Trp-Ile720Ala-Thr721Val-Asp722Asn substitutions in pT25-PorZ$_V$ | This study |
| pT25-PorZ$_V$-D* | Tyr755Ala-Val757Trp-Val759Trp substitutions in pT25-PorZ$_V$ | This study |
| pT25-PorZ$_V$-E* | Lys768Ala-Arg771Ala substitutions in pT25-PorZ$_V$ | This study |
| pTolB-T25 | *E. coli tolB* without signal sequence, cloned upstream T25 into pKT25 | Battesti and Bouveret, 2007 |

### OLIGONUCLEOTIDES

| **Oligonucleotide** | **Sequence (5' to 3')** |
|---|---|
| Expression vectors[a,b] | |
| 5-pIBA4-VSVG-RgpB-CTD | GTTTCGCTACCGTAGCGCAGGCCGCT*TATACAGATATTGAAATGAATAGATTAGGAAAA*GGTACATCTATTGCCGACGTAGCCAATG |
| 3-pIBA4-RgpB-CTD | GCCTTTTTCGAACTGCGGGTGGCTCCAGCTTTACTTCACTATAACCTTTTCTGTATACGTCTTGCC |
| 5-pIBA4-VSVG-Hpb35-CTD | GTTTCGCTACCGTAGCGCAGGCCGCT*TATACAGATATTGAAATGAATAGATTAGGAAAA*GGGCAAGAAAGTCTTGATAAAGCAGAGCC |
| 3-pIBA4-Hpb35-CTD | GCCTTTTTCGAACTGCGGGTGGCTCCAGCTTCAAGGAACTAAGACTTTAAGGAAATGCATTACACC |
| 5-pIBA4-VSVG-PPAD-CTD | GTTTCGCTACCGTAGCGCAGGCCGCT*TATACAGATATTGAAATGAATAGATTAGGAAAA*CTTCGTGCATGGTTCAACGCCGG |

| | |
|---|---|
| 3-pIBA4-PPAD-CTD | GCCTTTTTCGAACTGCGGGTGGCTCCAGCTTTATTTGAGAATTTTCATTGTCTCACGG |
| 5-pIBA4-VSVG-PorU-CTD | GTTTCGCTACCGTAGCGCAGGCCGCT*TATACAGATATTGAAATGAATAGATTAGGAAAA*TCATTCAGAGTGGTAGAT GGCATTGCTCC |
| 3-pIBA4-PorU-CTD | GCCTTTTTCGAACTGCGGGTGGCTCCAGCTCTATTGTGCTACCACGATCATTTTCTTGGCC |
| 5-pIBA4-VSVG-PorZ-CTD | GTTTCGCTACCGTAGCGCAGGCCGCT*TATACAGATATTGAAATGAATAGATTAGGAAAA*GGTACGGGGAGTGGATCA GCTTCC |
| 3-pIBA4-PorZ-CTD | GCCTTTTTCGAACTGCGGGTGGCTCCAGCTTCAGCGAATCACTGCGAAGCGAATTAG |

Bacterial Two-Hybrid (BACTH) [a,c]

| | |
|---|---|
| 5-BACTH-RgpB-CTD | GAAGtctagaTGGTACATCTATTGCCGACGTAGCCAATG |
| 3-BACTH-RgpB-CTD | GAAGggtaccCCCTTCACTATAACCTTTTCTGTATACGTCTTGCC |
| 5-BACTH-Hpb35-CTD | GAAGtctagaTGGGCAAGAAAGTCTTGATAAAGCAGAGCC |
| 3-BACTH-Hpb35-CTD | GAAGggtaccCCAGGAACTAAGACTTTAAGGAAATGCATTACACC |
| T25N-5-PpaD-CTD | GGCGGGCTGCAGATTATAAAGATGACGATGACAAGCTTCGTGCATGGTTCAACGCCGG |
| T25T18N-3-PPAD-CTD | CGAGGTCGACGGTATCGATAAGCTTGATATCGAATTCTAGTTATTTGAGAATTTTCATTGTCTCACGG |
| T25N-5-PorU-CTD | GGCGGGCTGCAGATTATAAAGATGACGATGACAAGTCATTCAGAGTGGTAGATGGCATTGCTCC |
| T25T18N-3-PorU-CTD | CGAGGTCGACGGTATCGATAAGCTTGATATCGAATTCTAGCTATTGTGCTACCACGATCATTTTCTTGGCC |
| T25N-5-PorZ-CTD | GGCGGGCTGCAGATTATAAAGATGACGATGACAAGGGTACGGGGAGTGGATCAGCTTCC |
| T25T18N-3-PorZ-CTD | CGAGGTCGACGGTATCGATAAGCTTGATATCGAATTCTAGTCAGCGAATCACTGCGAAGCGAATTAG |

Site-directed mutagenesis [d,e]

| | |
|---|---|
| A-RgpB-CTD-Bmotif | GAAAGTCCTGCTGCCGGGtggACGgccgtcaatATGAACGGCCGTCGTG |
| B-RgpB-CTD-Bmotif | CACGACGGCCGTTCATattgacggcCGTccaCCCGGCAGCAGGACTTTC |
| A-RgpB-CTD-Dmotif | CAAAACGGCGTGgctGCCtggCGCtggGCTACTGAAGGCAAGAC |
| B-RgpB-CTD-Dmotif | GTCTTGCCTTCAGTAGCccaGCGccaGGCagcCACGCCGTTTTG |
| A-RgpB-CTD-Emotif-IBA | GCAAGACGTATACAGAAgcgGTTATAGTGgcgTAAAGCTGGAGCCACCC |
| B-RgpB-CTD-Emotif-IBA | GGGTGGCTCCAGCTTTAcgcCACTATAACcgcTTCTGTATACGTCTTGC |
| A-RgpB-CTD-Emotif-T25 | GCAAGACGTATACAGAAgcgGTTATAGTGgcgGGGGTACCTAAGTAAC |
| B-RgpB-CTD-Emotif-T25 | GTTACTTAGGTACCCCcgcCACTATAACcgcTTCTGTATACGTCTTGC |

| | |
|---|---|
| A-PpaD-CTD-Bmotif | CGCCGGCACATATCGGtggAAGgctgttgacACCGCAGGAGAAGAAGTC |
| B-PpaD-CTD-Bmotif | GACTTCTTCTCCTGCGGTgtcaacagcCTTccaCCGATATGTGCCGGCG |
| A-PpaD-CTD-Dmotif | CTCCGGGCACAgctGTTtggGTTtggGAAGGAAATGGAATCCG |
| B-PpaD-CTD-Dmotif | CGGATTCCATTTCCTTCccaAACccaAACagcTGTGCCCGGAG |
| A-PpaD-CTD-Emotif-IBA | GAATCCGTGAGACAATGgcaATTCTCgcaTAAAGCTGGAGCCACC |
| B-PpaD-CTD-Emotif-IBA | GGTGGCTCCAGCTTTAtgcGAGAATtgcCATTGTCTCACGGATTC |
| A-PpaD-CTD-Emotif-T25 | GAATCCGTGAGACAATGgcaATTCTCgcaTAACTAGAATTCGATATC |
| B-PpaD-CTD-Emotif-T25 | GATATCGAATTCTAGTTAtgcGAGAATtgcCATTGTCTCACGGATTC |
| A-PorZ-CTD-Bmotif | CTGCAAGCCGGCTGTAGTtggAAAgccgtcaatACCACCGGCAGACTGC |
| B-PorZ-CTD-Bmotif | GCAGTCTGCCGGTGGTattgacggcTTTccaACTACAGCCGGCTTGCAG |
| A-PorZ-CTD-Dmotif | CTTCGGGCGTAgctGCCtggGCAtggTACGATCCGGTATCGAA |
| B-PorZ-CTD-Dmotif | TTCGATACCGGATCGTAccaTGCccaGGCagcTACGCCCGAAG |
| A-PorZ-CTD-Emotif | CGGTATCGAAAAAGTCCgcaCTAATTgccTTCGCAGTGATTCGC |
| B-PorZ-CTD-Emotif | GCGAATCACTGCGAAggcAATTAGtgcGGACTTTTTCGATACCG |

[a] sequence annealing on the target vector underlined
[b] VSVG tag coding sequence italicized
[e] restriction site in lower case (XbaI at the 5’ end, TCTAGA ; KpnI at the 3’ end, GGTACC)
[d] mutagenized codons in lower case
[e] base substitutions underlined
